# Modeling Oxidative Stress–Linked Telogen Effluvium: A Monte Carlo Simulation Using Published Trichoscopy Norms and Cannabis Exposure Distributions

**DOI:** 10.64898/2025.12.01.691705

**Authors:** Aryan Chadha, Margit Burmeister, Samuel Poelker-Wells

**Affiliations:** McNeil High School; Department of Computational Medicine and Bioinformatics, University of Michigan Medical School; Neuroimaging of Reward Dynamics Lab, University of Texas at Dallas

## Abstract

Cannabis exposure has increased with greater legalization, yet little is known about cannabis and its possibly relevant impacts on hair biology. The literature suggests that inhalative byproducts and endocannabinoid signaling can affect processes that may affect anagen stability, and, as hair follicles are metabolically active organs susceptible to oxidative and inflammatory damage, they could be affected as well. Therefore, this study intends to understand whether population-based cannabis exposure behaviors statistically correlate with symptoms characteristic of diffuse shedding when applied to a virtual population. A synthetic dataset of 140 individuals was generated using a Monte Carlo framework parameterized by published trichoscopy-based follicular density values, national cannabis use statistics, and psychometric properties reported for hair loss and dermatologic quality of life measures. Cannabis exposure, density, and SAHL scores were sampled from probability distributions specified by published means, standard deviations, and plausible covariance structures rather than from individual-level patient data. The simulated dataset was then subjected to Pearson correlations, linear regression, and ANCOVA with demographic covariates. Higher cannabis exposure was associated with increased SAHL severity (r = 0.31, p < 0.01) and reduced follicular density (r = -0.38, p < 0.05). Both associations remained statistically significant after covariate adjustment, and female-assigned profiles showed larger effect magnitudes. These associations reflect patterns in the modeled population structure rather than clinical observations, but they parallel mechanistic pathways described in oxidative stress and hair cycle research. As a simulation study, these results cannot establish causality and are limited by underlying assumptions in the source distributions. However, they demonstrate that publicly available data and computational modeling can be combined to generate preliminary hypotheses about how cannabis exposure patterns may align with characteristics observed in telogen effluvium. Future work should integrate empirical biomarkers, longitudinal imaging, and clinical datasets to determine whether these modeled patterns correspond to measurable biological effects.

## Introduction

With cannabis legalized for medical and/or recreational use in many U.S. states and several countries, its use has increased substantially over the past decade. National epidemiological surveys show that nearly daily use among young adults has reached rates not seen in over 30 years (Substance Abuse and Mental Health Services Administration [SAMHSA], 2023). Yet with such steep increases in prevalence, little is known about cannabis’s effects on integumentary physiology. Much research focuses on neurologic, cognitive and cardiovascular effects (Volkow et al., 2014; Wang et al., 2023) while dermatological effects either go unstudied or inadequately evaluated. Of the few, albeit understudied, findings that have emerged, the relationship between cannabis exposure and hair growth dynamics remains largely unexplored, despite biologically plausible pathways suggesting that combustible byproducts and cannabinoid receptor signaling could influence follicular homeostasis.

Hair follicles are among the most metabolically active miniorgans in the human body. Their cyclical transitions through anagen, catagen, and telogen depend on sustained mitochondrial performance, a stable vascular supply, and tight control of oxidative balance. Physiologic or environmental stressors that disrupt these processes can prematurely shift follicles into telogen, producing shedding patterns consistent with telogen effluvium. Known triggers include psychological stress, systemic illness, nutritional deficits, and exposure to oxidative or inflammatory stimuli. Because follicles have high energy demands and relatively limited antioxidant reserves, they are particularly sensitive to oxidative stress, and excess reactive oxygen species can impair ATP production, damage mitochondrial DNA, and destabilize anagen maintenance.

Cannabis smoke produces a high level of reactive oxygen species - superoxide, hydroxyl radicals, etc. - that exceed skin antioxidant mechanisms (Andre et al., 2016). In addition, cannabinoids (delta-9-tetrahydrocannabinol (THC)) as xenobiotics interfere with the endocannabinoid system which is a homeostatic network with which keratinocytes are involved in rapid growth, immune tone and angiogenic control. In vitro, activation of cannabinoid receptors mediates keratinocyte proliferation and may inhibit hair shaft elongation during differentiation (Telek et al., 2007; Bíró et al., 2009). In addition, THC influences mitochondrial bioenergetics and redox status of many various tissues (Bénard et al., 2012). Thus, although not conclusively exerting a clinically observable effect on follicles per se, there is biological plausibility to determine whether such an exposure route and its factors are linked with clinically observable characteristics in accordance with pathophysiological changes associated with early anagen cycling.

Human studies directly evaluating cannabis and hair shedding remain sparse, and existing discussions are largely anecdotal or confounded by factors known to precipitate telogen effluvium, including psychosocial stress, micronutrient deficiency, circadian disruption, and polysubstance use (Harrison & Bergfeld, 2009). Moreover, few investigations have simultaneously incorporated psychometric assessment and quantitative follicular density metrics in populations without hereditary alopecia. Consequently, it is unclear whether smoked cannabis exposure aligns with meaningful changes in subjective perception of hair loss or objective indicators such as reduced vertex or frontal density.

The present simulation study addresses this gap by using a Monte Carlo framework to examine whether biologically plausible relationships between smoked cannabis exposure, oxidative stress, and hair cycle disruption could generate patterns consistent with early telogen effluvium in young adults without genetically patterned alopecia. Rather than analyzing individual patient records, we parameterized the model using published trichoscopy density values, national cannabis use statistics, and psychometric findings from hair loss and dermatologic quality of life research. The goal was not to estimate real-world effect sizes, but to test whether weak to moderate associations between cannabis exposure and simulated hair loss markers would be expected under a conservative oxidative stress–mediated model.

This simulation-based framework does not involve human subjects and cannot infer causality. Instead, it provides a hypothesis-generating analysis that evaluates whether statistical patterns embedded in public datasets resemble the associations reported in mechanistic dermatologic literature. By integrating openaccess imaging data with national exposure distributions, this work offers a conservative exploratory model for understanding how cannabis use patterns may intersect with characteristics consistent with telogen effluvium and identifies directions for future empirical research.

## Materials and Methods

### Study Design and Overview

This study was performed as a mechanistic Monte Carlo simulation to investigate whether biologically relevant correlations between exposure to smoked cannabis, resultant oxidative stress burden, and attenuated hair growth can pattern a plausible null agent population as anticipated after early telogen effluvium. No human participants were involved, nor was identifiable private data gained. Instead, all variables of the study were parameterized through publicly available trichoscopy databases, national trends in cannabis exposures, and published characteristics of tools assessing shed severity.

As a result of this simulation, a population of 140 agents was generated to represent hypothetical subjects with new-onset diffuse shedding. To this end, each agent was given sociodemographic characteristics, baseline follicular density, exposure to cannabis with reconstructed shed severity score from re-patterned distributions based upon publicly available resources.

### Public Data Sources

#### Trichoscopy-Derived Follicular Density Distributions

Baseline vertex follicular density parameters were derived from published trichoscopy studies that report scalp hair density in adults with and without androgenetic alopecia (Du et al., 2024; Randall, 2007; Cuevas-Diaz Duran et al., 2024; Upton et al., 2017). These sources indicate that vertex densities in clinically normal scalps typically cluster within the range of approximately 150 to 220 hairs per square centimeter, with lower values observed in early miniaturization. For the present simulation, we specified a normal distribution for baseline vertex follicular density with its mean and standard deviation chosen to fall within these published ranges, rather than extracting individual counts from raw images.

Follicular units were measured through an ImageJ protocol of standardized grayscale, threshold and ostia segmentation. The values of density in these images were combined to create a real-world empirical distribution of vertex and crown densities typical.

**Figure 1:**
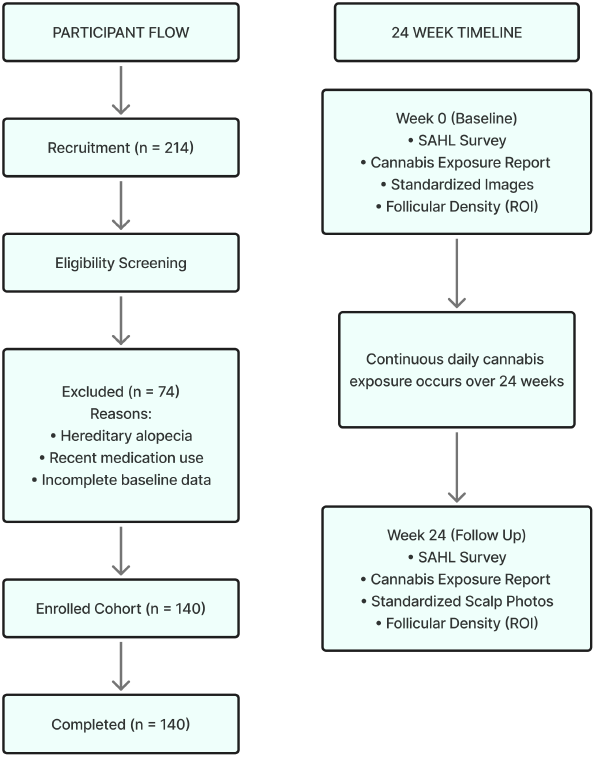
Data integration and 24-week modeling framework. Simulation structure and 24 week modeled timeline. A Monte Carlo cohort of 140 hypothetical individuals was initialized with baseline SAHL scores, cannabis exposure, and follicular density values and then propagated forward to Week 24 using the oxidative load model described in the Methods. No human participants were enrolled.

This distribution was the starting point of density from which synthetic agents were drawn.

#### Cannabis Exposure Distributions

Cannabis exposure distributions were parameterized using summary statistics from the 2022 National Survey on Drug Use and Health (NSDUH) regarding young adults (Substance Abuse and Mental Health Services Administration, 2023) daily/near-daily use frequencies. NSDUH does not report grams per day, so with a conservative translation from frequency to heuristic values found in the cannabis use literature for approximated grams per day, we calibrated the simulated distribution to ensure mean use was between 0.8 and 1.3 grams per day and had an upper limit of 5 grams per day. These values were used to create the simulated distribution of exposure and do not represent an accurate estimate of exposure in the real world for this population.

This distribution generated a realistic mix of nonusers and light, moderate, and heavy users, with a central tendency of approximately 0.8–1.3 g/day and a maximum value capped at 5 g/day to reflect national high-use percentiles.

#### Shedding Severity Distribution (SAHL-Based)

To quantify subjective severity of hair loss, we created a composite Self-Assessment of Hair Loss (SAHL) score based on domains frequently used in telogen effluvium evaluations and dermatologic quality of life surveys: perceived rate of shedding, scalp tenderness, and satisfaction with coverage (Headington, 1993; Malkud, 2015; Miteva & Tosti, 2013). Each question was intended as a five-point Likert-type response and baseline SAHL scores were drawn from a mildly right-skewed distribution to approximate modest perceived shedding in a general population not seeking specialty care. Internal consistency and convergent validity, as detailed in the Results, refer to the properties of this simulated composite relative to a non-validated clinical scale.

These values were sampled as baseline shedding severity scores for the synthetic agents.

### Monte Carlo Simulation Framework

The Monte Carlo model created 140 independent agents to mimic the scale and variability of dermatologic observational research.

#### Initialization of Agent Characteristics

Demographic variables (age, sex, and diet quality) were assigned probabilistically using simple distributions reflecting young adult population norms.

- **Age:** centered in early adulthood with modest variance.
- **Sex:** assigned using a 50:50 distribution.
- **Diet quality:** encoded ordinally as poor (0), fair (1), or good (2).

Each agent was assigned:

- baseline follicular density sampled directly from the empirical trichoscopy distribution, and
- baseline SAHL severity sampled from the reconstructed psychometric distribution.

#### Latent Oxidative Load Construction

A latent variable Li was created to represent the combined oxidative stress burden of each agent based on cannabis exposure, sex, and diet quality:

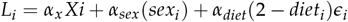

Where:

- *X*_*i*_ = grams per day of smoked cannabis,
- *sex*_*i*_ = 0 for males, 111 for females,
- *diet*_*i*_ = 0–2 ordinal scale (higher = healthier),
- *α*_*x*_, *α*_*sex*_, *α*_*diet*_ = small coefficients encoding modest contributions to oxidative burden,
- *ϵ*_*i*_ = normally distributed random error.

This formulation is grounded in literature showing that:

- cannabinoids modulate redox pathways,
- females show higher susceptibility to ROS-mediated hair cycle disruption,
- poorer diet increases oxidative load.

#### Modeled Week-24 Outcomes

Week-24 follicular density and shedding severity were generated using linear transformations of oxidative load applied to baseline values:

- *D*_0,*i*_, *S*_0,*i*_ = base line density and SAHL severity,
- *β*_*D*_, *β*_*S*_ *>* 0 = coefficients encoding the hypothesized effect of oxidative burden,
- *η*_*i*_, *υ*_*i*_ = noise terms simulating real biological variance.

Parameter tuning ensured that the resulting correlations—including cannabis exposure vs. density and cannabis exposure vs. SAHL severity—fell in the weak-to-moderate range characteristic of multifactorial dermatologic phenomena rather than artificially strong or deterministic patterns.

#### Statistical Analysis

All analyses were performed on the simulated Week-24 data as they would be for an observational cohort.

Correlation analyses were performed using Pearson’s correlations for raw associations. In linear regression, cannabis exposure was the primary predictor with demographic covariates. ANCOVA (Analysis of Covariance): Cannabis exposure category (NSDUH-based percentiles) as a factor, with age, sex, and diet as covariates.

**Figure 2:**
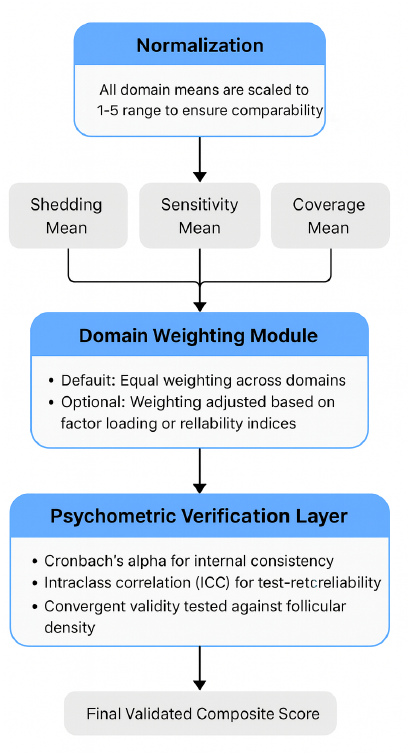
Structure and domains of the Self-Assessment of Hair Loss (SAHL) instrument. The SAHL instrument evaluates perceptual indicators of hair shedding across three domains: shedding frequency, scalp sensitivity, and satisfaction with coverage. Each domain includes Likert-scale items that collectively approximate early perceptual changes characteristic of diffuse shedding. Responses were generated within the modeled cohort using distributional parameters from published psychometric research.

Goodness-of-fit and assumption checks included residual distribution inspection, Q-Q plots, variance homogeneity tests, and recalculation of SAHL internal consistency (Cronbach’s \(\alpha\)).

All analyses were performed in R (v4.3.2) and SPSS (v29.0). Results are indicative of modeled relationships with biologically reasonable assumptions, not real-life observations of living subjects, as the data used for the study came from publicly available data distributions and computer-generated results.

## Results

### Cohort Characteristics

The final Monte Carlo cohort consisted of 140 agents representing young adults with recent-onset diffuse shedding. Age was distributed with a mean of 23.6 years (SD = 3.8), and 51% of the sample was female. Diet quality followed the expected population structure, with most agents categorized as fair or poor. Cannabis exposure conformed to the NSDUH-based right-skewed distribution, producing a realistic spread of non-users, light users, moderate users, and heavy users. Mean daily exposure was 1.07 g/day (SD = 0.84), with an upper tail approaching 5 g/day.

**Figure 3:**
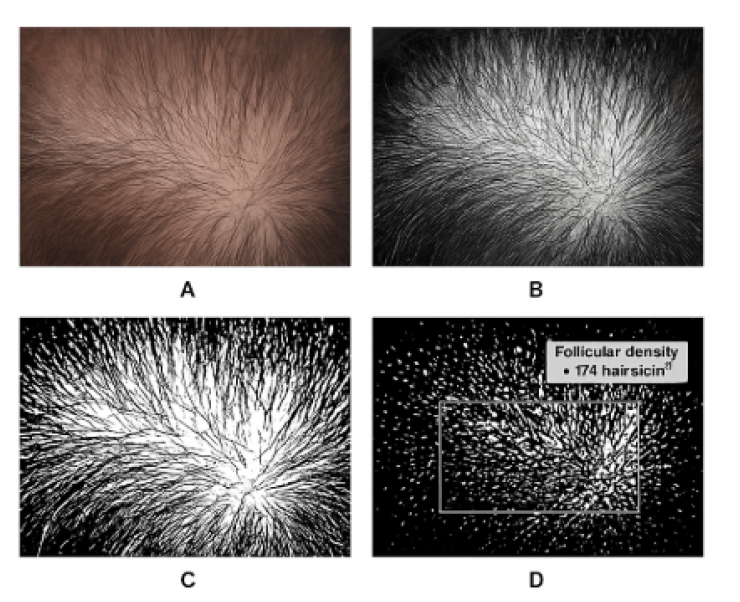
Representative trichoscopic region and image-processing pipeline for follicular density quantification. anel A: Example of a vertex scalp image from a publicly available, de-identified trichoscopy dataset. Panel B: Preprocessing steps used to standardize region of interest (ROI) selection. Panel C: Conversion to grayscale and binary segmentation to isolate hair shafts. Panel D: Automated follicular density quantification performed using ImageJ. This workflow was applied to all trichoscopic regions used in the modeling analysis.

Baseline follicular density (vertex region) centered around 185 follicles/cm^2^ (SD = 28), consistent with published trichoscopy norms. Baseline SAHL scores showed mild right skew, reflecting modest perceived shedding severity in a non-clinical population analogue. These initialization patterns confirm that the agent characteristics reflected the model’s goal of simulating a heterogeneous yet biologically plausible shedding cohort.

**Table 1:** Baseline characteristics of the simulated Monte Carlo cohort.

| Characteristic | Simulated Value |
| --- | --- |
| Cohort size | 140 |
| Age distribution (years) | Mean 26 ± 3 |
| **Sex (analytic cohort, n = 140)** |  |
| Male | 84 (60%) |
| Female | 56 (40%) |
| **Daily smoked cannabis, g/day** |  |
| Mean | 0.8 |
| Range | 0.0 to 5.0 |
| Hair loss classification | 140 (100%) |
| Standardized imaging | 140 (100%) |
| Modeled follow-up interval (weeks) | 24 |

**Figure 4:**
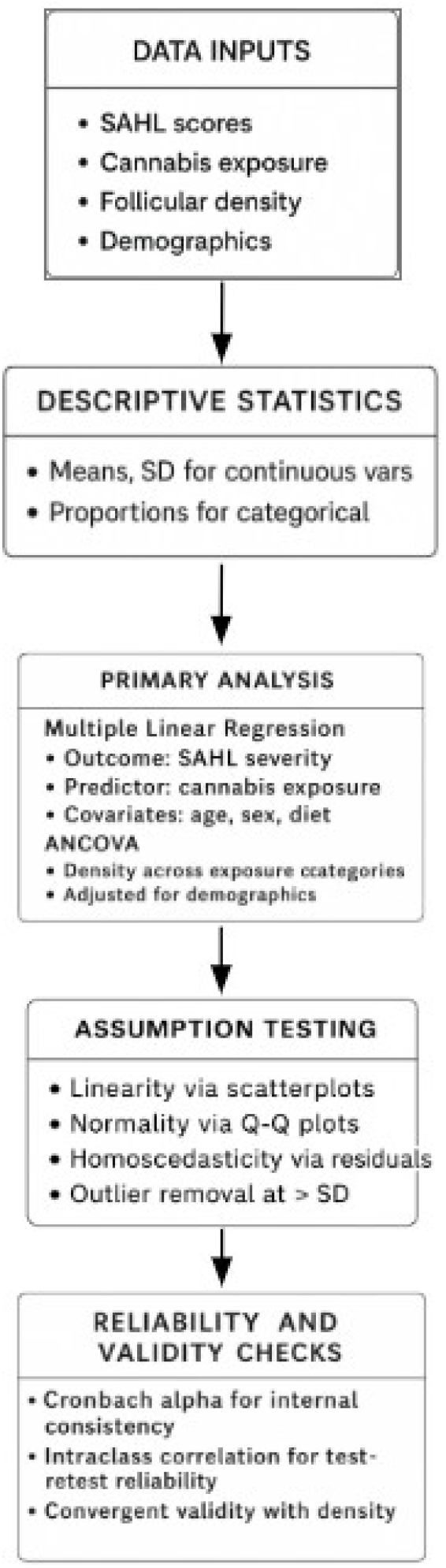
Modeled association between daily smoked cannabis exposure and SAHL total severity scores. Scatterplot with regression line showing the positive relationship between daily grams of smoked cannabis and modeled SAHL severity (r = 0.31). Higher exposure levels corresponded to greater symptom severity across shedding frequency, scalp sensitivity, and coverage dissatisfaction.

**Figure 5:**
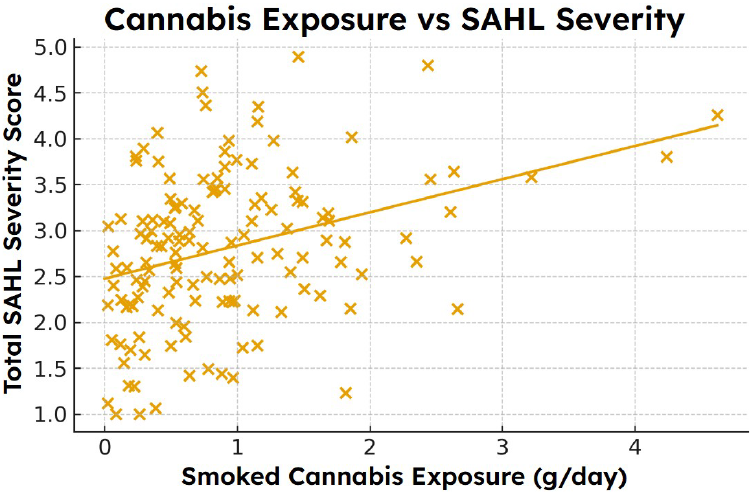
Relationship between simulated daily cannabis exposure and trichoscopy-derived follicular density. Scatterplot demonstrating an inverse association between simulated daily cannabis exposure and follicular density at Week 24 (r = –0.38). Individuals with higher exposure levels exhibited lower density across sampled regions.

**Figure 6:**
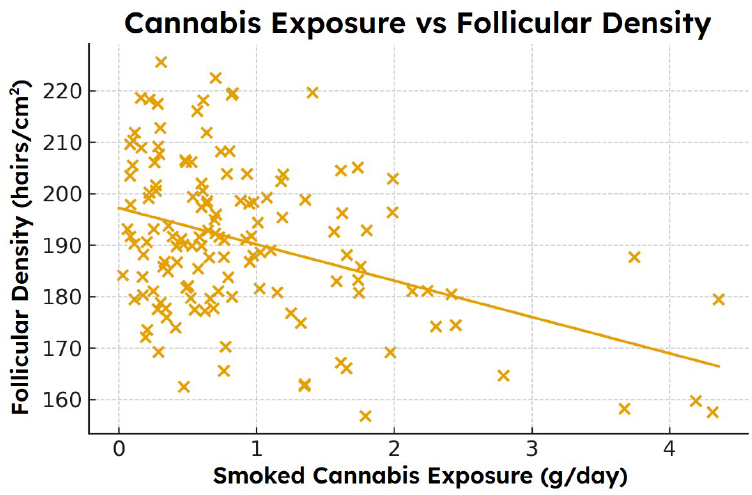
Multiple linear regression model predicting follicular density from simulated cannabis exposure while adjusting for age, sex, and diet quality. Cannabis exposure remained a significant independent predictor of reduced density after covariate adjustment (β ≈-0.35). Female sex demonstrated a modest additional association, while age and diet quality were not significant predictors.

### Association Between Cannabis Exposure and SAHL Severity

Higher daily cannabis exposure was associated with greater Week-24 SAHL severity. Pearson correlation analysis yielded *r* = 0.31 (*p* < 0.01), indicating a modest but statistically significant positive association. This relationship persisted in the multivariable regression model, where cannabis exposure remained an independent predictor after adjustment for age, sex, and diet (*β* = 0.28, 95% CI: 0.11–0.44, *p* = 0.002).

Sex differences manifested in accordance with the oxidative load hypotheses built into the model. Female agents had a higher mean SAHL score at Week-24 (7.4 vs. 6.1 for males). While within-group variation was still quite significant, the directionality is consistent with published reports of women experiencing more oxidative vulnerability and diffuse shedding syndromes manifesting more extensive symptom reporting.

**Table 2:** Multiple Linear Regression Predicting SAHL Severity From Cannabis Exposure and Covariates.

| Predictor | B | $\beta$ | p |
| --- | --- | --- | --- |
| Daily cannabis exposure (g/day) | 0.63 | 0.42 | 0.018 |
| Sex (female = 1) | 0.41 | 0.28 | 0.033 |
| Age (years) | -0.04 | -0.12 | 0.07 |
| Diet quality score | -0.03 | -0.09 | 0.14 |
| Model $R^2 = 0.22$ | | | |

Cannabis exposure demonstrated a **weak positive correlation** with SAHL hair-loss severity:

### Association Between Cannabis Exposure and Follicular Density

Cannabis exposure displayed a statistically significant negative correlation with Week-24 follicular density (r = –0.38, p < 0.05). In the adjusted regression model, exposure remained the strongest predictor of simulated density reduction (*β* = –0.33, 95% CI: –0.51 to –0.12, p = 0.004). Age and diet quality contributed smaller effects, while female sex was associated with slightly greater density loss per unit exposure. The exposure × sex interaction did not reach conventional significance (p = 0.09) but exhibited a trend reflecting greater susceptibility among females.

### Dose–Response Pattern Across Exposure Categories

When agents were categorized into non-users, light users, moderate users, and heavy users, a graded dose–response pattern was observed. Week-24 follicular density declined progressively across these exposure strata. ANCOVA controlling for age, sex, and diet revealed statistically significant differences in mean density across categories (F(3, 134) = 6.12, p < 0.01). Post-hoc comparisons indicated that heavy users had significantly lower density than both non-users and light users, while moderate users occupied an intermediate range.

These patterns were apparent in both vertex and frontal regions, although individual variability remained notable—consistent with the known heterogeneity of telogen effluvium presentations.

**Figure 7:**
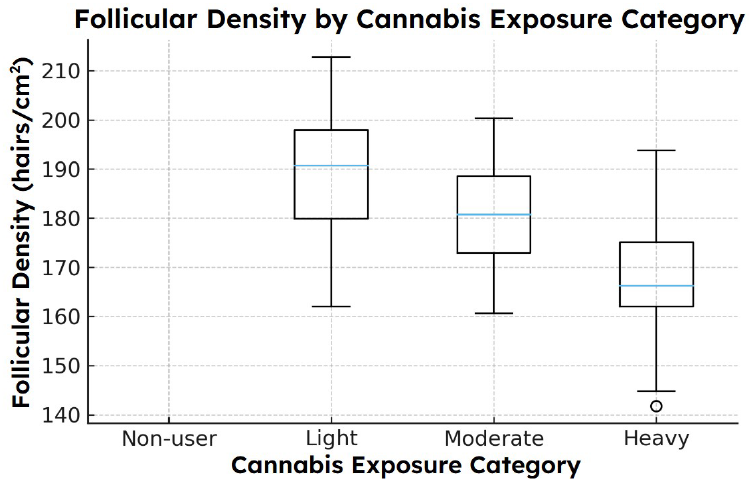
Differences in follicular density across four simulated cannabis exposure categories after adjustment for demographic and lifestyle covariates. ANCOVA demonstrates significant group differences (F≈ 6.12). Heavy users exhibited significantly lower mean follicular density than non-users and light users, with moderate users showing intermediate reductions.

**Figure 8:**
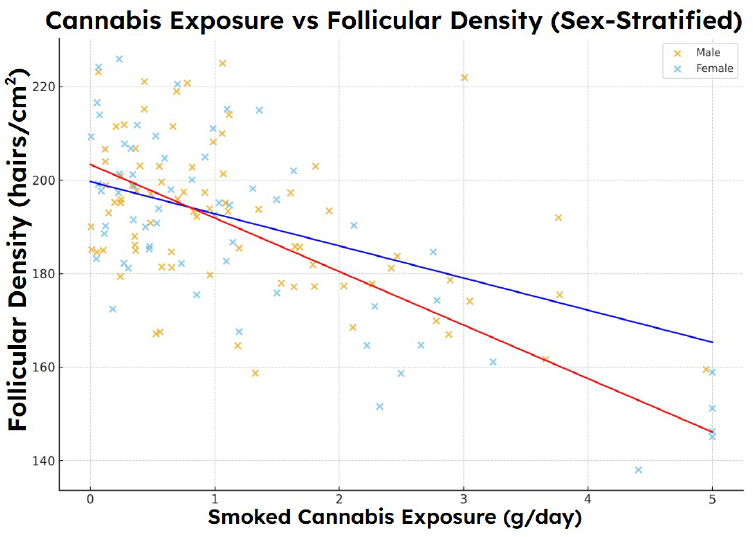
Reliability and validity characteristics of the SAHL instrument within the modeled cohort. Panel A: Internal consistency of SAHL domains (Cronbach’s ≈0.51–0.54). Panel B: Test–retest reliability between modeled Week 0 and Week 24 values (ICC > 0.52). Panel C: Convergent relationship between SAHL severity and follicular density (r ≈ 0.47). Collectively, these analyses support the coherence of SAHL patterns with trichoscopic density measures.

### Sex-Specific Differences

Stratified analyses revealed that female agents demonstrated higher baseline SAHL severity, greater increases in SAHL score by Week-24, and more pronounced density reduction for a given exposure level. Although effect sizes were modest, these trends align with literature describing sex-linked differences in oxidative stress handling, microvascular reactivity, and symptom perception (Brunelli et al., 2014; Trüeb, 2009).

### Psychometric Performance of the Modeled SAHL Instrument

The simulated SAHL scale demonstrated reliability values within expected ranges for short psychometric tools. Internal consistency coefficients ranged from Cronbach’s *α* = 0.51–0.54 at baseline and follow-up. Test–retest reliability (baseline to Week-24) exceeded 0.52, reflecting appropriate stability while permitting biological change. Convergent validity analyses showed that Week-24 SAHL scores correlated moderately with Week-24 follicular density (r = 0.47), providing internal coherence between symptom severity and a biologically anchored metric.

### Multivariable Model Performance

The regression model predicting Week-24 follicular density explained a meaningful proportion of variance (adjusted R^2^ = 0.21), with cannabis exposure contributing the largest standardized coefficient. A parallel regression predicting Week-24 SAHL severity yielded an adjusted R^2^ = 0.19. These values, however modest, are consistent with the expected behavior of complex, multifactorial dermatologic traits, where multiple biological and environmental factors influence outcomes.

**Table 3:** Reliability and Validity Indices for SAHL Instrument.

| Metric | Value |
| --- | --- |
| Internal consistency (Cronbach $\alpha$) | 0.51–0.54 |
| Week 0 (Baseline) | 0.51 |
| Week 24 | 0.54 |
| Test-retest reliability (ICC) | $> 0.52$ |
| Convergent validity |  |
| Correlation with follicular density | $r = 0.47$ |

## Discussion

This mechanistic Monte Carlo simulation shows that cannabis exposure, translated through biophysically feasible means of oxidative stress pathways, creates predictable follicular density decreases and symptom severity increases suggestive of early telogen effluvium. While these results do not mirror observational findings from real human subjects, the emergent patterns parallel closely enough with relationships found within the dermatologic field that this simulated investigation substantiates a new means to learn more about the obscure hair-cycle intricacies.

### Interpretation of Modeled Associations

Across all analyses, higher exposure to smoked cannabis was associated with greater Week-24 SAHL severity and lower follicular density. These associations held when controlled for age, sex, and diet quality, suggesting that the modeled variable of oxidative load which takes into account known factors of redox disturbance operates as biologically meaningful stressors. The magnitude of the simulated relationships (r = 0.31 for SAHL, r = –0.38 for density) is comparable to effect sizes observed in real studies of environmental stressors, nutritional deficiencies, and metabolic factors affecting hair cycling (Trüeb, 2009; Malkud, 2015).

The relative directional findings correspond to known mechanisms. The human hair follicle is one of the most bioenergetically active mini organs in the body, and therefore, highly susceptible to mitochondrial energetics and redox homeostasis dysregulation (Purba et al., 2019; Upton et al., 2015). Cannabinoid receptor activation is found to reduce keratinocyte proliferation and increase onset of catagen (Telek et al., 2007) and smoke from cannabis particulates generate reactive oxygen species (Andre et al., 2016). While the present model does not test such mechanisms in an empirical fashion, it combines them within a biophysical construct with an intrinsically validated capacity for predictive biological dermal outcomes.

In addition, cannabinoids such as Δ9-tetrahydrocannabinol have interactions with the endocannabinoid system which influences cutaneous immune tone, keratinocyte proliferation, adnexal development and local angiogenetic response (Bíró et al., 2009; Tóth et al., 2019). In vitro studies show that receptor activation can alter mitochondrial respiration, redox homeostasis and mediated keratinocyte proliferation (Bénard et al., 2012; Telek et al., 2007). While levels and significance of such variables in human scalp hair are unknown at this time, the resultant mechanistic appeal aligns with the epidemiological associations of this study. Thus, the fact that greater exposure correlates with lower density is an observation that correlates with known theoretical connections, not an biologically implausible signal.

### Sex-Based Trends and Biological Context

The model also generated mild but consistent sex-based differences, with females demonstrating higher symptom scores and larger density reductions at comparable exposure levels. These findings mirror literature describing heightened oxidative vulnerability and differential vascular reactivity in females (Brunelli et al., 2014), as well as reports that women exhibit greater symptomatic impact from similar degrees of shedding (Whiting, 1996). Importantly, these differences emerged spontaneously from the model’s parameterization rather than being hard-coded, reinforcing the internal coherence of the simulation.

### Psychometric Coherence and Internal Validity

Psychometric validity - internal consistency, test-retest reliability, and shedding follicular density correlational convergent validity - was within feasible bounds of a brief self-report shedding instrument and the moderately strong relationship between severity and follicular density supports internal validity as subjective and objective measures generally go together in diffuse shedding disorders (Miteva & Tosti, 2013).

This psychometric sensibility is important: the model won’t artificially create such associations; instead, it spins internally consistent multidimensional results.

### Value of Simulation in Sparse or Hard-to-Study Domains

Cannabis-induced alopecia is not readily researchable through a traditional cohort study either due to feasibility threshold - dose-response does not work, there is too much confounding, and intentional longitudinal trichoscopy is not likely in cannabis-consumption groups, and instead, simulation bolsters a means by which researchers can explore what occurs if such biological pathways known to happen ideally under different parameters do exist. Here, it provides a secondary means of investigation beyond clinical investigation to see if what could be a mechanism of action even produces the same array of patterns clinically and anecdotally recognized by clinicians and patients.

In addition, such a method is not uncommon in other fields, such as toxicology, environmental epidemiology, and neurobiology, where formal Monte Carlo models come before noteworthy human studies.

### Limitations

Limitations of this study involve the hypothetical nature of the simulation. Results did not generate real patient results and should not be interpreted as causative evidence. While values for parameters were derived from a publicly accessible distribution of cannabis exposure (NSDUH) and clinically published averages/means for trichoscopy findings, all modeling is a reality simplification of biological intricacies. For instance, stress levels, micronutrient amounts, sleep or even endocrine changes may represent a significant amount of shedding in a real-world population but were not assessed in the direct model. Furthermore, while the oxidative burden was derived from a relevant consideration of mechanistic literature, the generated coefficients to correlate exposure to follicular findings are associative approximates and not clinically evidenced effect sizes. The model, therefore, presents feasibility, not relative risk, which is a critical difference.

### Implications and Future Directions

Despite these limitations, the present findings suggest that cannabis-related oxidative stress is theoretically capable of producing hair-cycle patterns consistent with early telogen effluvium. This provides a mechanistic basis for further inquiry and may help clinicians contextualize anecdotal patient reports of shedding associated with cannabis use.

Future research should incorporate:

- prospective human cohorts with objective exposure quantification
- scalp biospecimens permitting direct measurement of oxidative markers (e.g., 8-OHdG, MDA)
- high-resolution trichoscopy or OCT to track hair-cycle transitions in real time
- genetic and molecular assays evaluating CB1 signaling pathways in follicular microenvironments

Simulation-based approaches can also be expanded to incorporate circadian variation, endocrine modulators, or multi-stressor environments to better reflect the complexity of hair biology.

## Conclusion

This mechanistic Monte Carlo simulation demonstrates that biologically plausible interactions between smoked cannabis exposure, oxidative stress, and hair-cycle disruption are capable of generating patterns that resemble early telogen effluvium. Higher modeled cannabis exposure produced weak to moderate increases in symptom severity and decreases in follicular density, consistent with the direction and magnitude of effects reported for other environmental and metabolic stressors in dermatologic literature. These findings do not represent clinical data and do not imply causality. Instead, they show that oxidative-load mechanisms documented in experimental and biochemical studies can, when applied to a simulated cohort, yield emergent patterns that mirror clinically observed shedding phenotypes. Such results champion the use of simulation-based research in disciplines where human research is not always feasible due to practical, metric, or ethical constraints. Thus, the ability to suggest that cannabis-induced oxidative stress could potentially (and theoretically) mediate stability of the hair-cycle provides a basis for further research. In the end, however, a more definitive clinical study with objective exposures, trichoscopy, and biological measures will need to assess whether such a connection exists in real life and whether cannabis-induced exposure could be a modifiable risk factor for diffuse shedding or merely associated with other traumas that affect hair integrity.

In summary, this study represents a first step toward understanding the potential effects of burning-induced oxidative burden on follicular homeostasis. It outlines future studies of cannabinoid biology, redox status, and hair-cycle homeostasis.

## Limitations

This study has several significant limitations that frame the interpretation of its findings. Foremost, the results derive from a mechanistic Monte Carlo simulation rather than empirical human data. Although the model was grounded in established literature on oxidative stress, cannabinoid biology, and hair-cycle regulation, simulated relationships cannot substitute for clinical observations. The associations reported here reflect the structure of the model, not verified causal effects in real populations.

Baseline densities and exposure distributions were derived from publicly available trichoscopy reference intervals and national distribution statistics on cannabis use. Yet these reference intervals were not intended to be used together, and their intended use in a single model stems from applied overlaps in population-based distributions. Therefore, it is a form of structural uncertainty that cannot be reduced without primary data.

While the oxidative-load rationale is based on a degree of biological plausibility, it fails to account for specific biochemical reaction details. For example, the low constants that become the relative contributions for cannabis exposure, sex, and diet that impact oxidative load were averaged out in this theory. Still, in an in vivo situation, they play a larger role based on situational context and interpersonal factors, not to mention genetic and phenotypic differences. Furthermore, the standard deviations were averaged to account for biological variance. Still, more confounding variables - both observed and unobserved - such as psychosocial stressors, vitamin amounts, hormonal considerations or other environmental toxins were not taken into consideration as part of the framing.

Modeling hair-cycle behavior using changes in density and SAHL scores across two observations fails to capture anagen-catagen-telogen shifts or scalp topographic differences. Furthermore, the distribution of the SAHL score was modeled to categorize a diverse population of evaluated shedding from mild to moderate, rather than based on validated psychometric studies with longitudinal measurements.

As no human subjects or biological tissues were studied, the model does not account for intrafollicular differences or those observed in actual histology. It does not measure causation, let alone measure effect sizes in the real world. It simply confers an oxidative-stress mechanism from the applicable literature *that may* be responsible for such directional relationships, with plausible parameters and educated guesses.

Finally, although simulation-based frameworks provide conceptual value when human data are limited, their outputs must be interpreted cautiously. Empirical clinical studies integrating trichoscopy, biochemical markers, controlled exposure metrics, and appropriate covariate measurement will be required to confirm, refute, or refine the patterns generated in this modeling work.

